# Clustering co-abundant genes identifies components of the gut microbiome that are reproducibly associated with colorectal cancer and inflammatory bowel disease

**DOI:** 10.1101/567818

**Authors:** Samuel S. Minot, Amy D. Willis

**Affiliations:** Microbiome Research Initiative, Fred Hutchinson Cancer Research Center, Seattle, Washington, USA; Department of Biostatistics, University of Washington, Seattle, Washington, USA

## Abstract

**Background:** Whole-genome “shotgun” (WGS) metagenomic sequencing is an increasingly widely used tool for analyzing the metagenomic content of microbiome samples. While WGS data contains gene-level information, it can be challenging to analyze the millions of microbial genes which are typically found in microbiome experiments. To mitigate the ultrahigh dimensionality challenge of gene-level metagenomics, it has been proposed to cluster genes by co-abundance to form Co-Abundant Gene groups (CAGs). However, exhaustive co-abundance clustering of millions of microbial genes across thousands of biological samples has previously been intractable purely due to the computational challenge of performing trillions of pairwise comparisons.

**Results:** Here we present a novel computational approach to the analysis of WGS datasets in which microbial gene groups are the fundamental unit of analysis. We use the Approximate Nearest Neighbor heuristic for near-exhaustive average linkage clustering to group millions of genes by co-abundance. This results in thousands of high-quality CAGs representing complete and partial microbial genomes. We applied this method to publicly available WGS microbiome surveys and found that the resulting microbial CAGs associated with inflammatory bowel disease (IBD) and colorectal cancer (CRC) were highly reproducible and could be validated independently using multiple independent cohorts.

**Conclusions:** This powerful approach to gene-level metagenomics provides a powerful path forward for identifying the biological links between the microbiome and human health. By proposing a new computational approach for handling high dimensional metagenomics data, we identified specific microbial gene groups that are associated with disease that can be used to identify strains of interest for further preclinical and mechanistic experimentation.

## Background

Metagenomic analysis of the microbiome typically falls into the categories of taxonomic classification, metabolic pathway reconstruction, or genome reconstruction. While each has been used to good effect, each also has its own limitations. Taxonomic analysis is constrained by the size and quality of reference databases, which have started to provide decreasing taxonomic precision as the number of sequenced genomes grows [1]. Metabolic analysis is limited by our ability to annotate biochemical function from primary sequence, with only a minority of genes receiving any sort of annotation. Genome reconstruction (or “genome-resolved metagenomics”) has made immense contributions to our understanding of microbial diversity and evolution, but is challenging to exhaustively characterize environments like the human gut, which contain hundreds or thousands of strains. In contrast, we took the approach of quantifying each individual gene *de novo* from a given metagenome. While this approach presented considerable computational challenges, it is unconstrained by the limitations of reference databases or annotation systems and therefore presents the possibility of discovering novel biological patterns in the human microbiome.

While the microbiome has been implicated in a number of human diseases, we chose to focus on CRC and IBD because of the availability of metagenomic data from multiple independent cohorts[2–8]. Associative studies characterizing differences in the microbiome as a function of disease status are complicated by the effect of disease and treatment process on the microbiome [9–12] but it is still possible that some of the differences in the microbiome may play some causal role or implicate a causal biological process.

The approach of gene-level metagenomics is not new to this study and has been proposed previously as an alternative to taxonomic or metabolic pathway analysis [13]. Indeed even the popular HUMAnN2 tool [14] includes gene-family abundance estimation using the UniRef database of proteins [15]. We took the previously-described approach of grouping together genes that are consistently found at a similar level of abundance across multiple samples [13]. Such co-abundant genes are likely to be found on the same chromosome or piece of DNA across multiple samples, such as in the core genome for a bacterial species or consortium, on a plasmid that may move between strains, or as part of an operon in the accessory genome of a species that is only found in a subset of strains. Biologically speaking, co-abundant genes are not independent entities, and can be grouped together for purposes of inferring their relationship with human health and disease. In addition, grouping genes by co-abundance finds low-dimensional structure in high-dimensional gene-level data, mitigating challenges with the statistical analysis of high-dimensional metagenomics data.

## Results and Discussion

The primary analytical challenge that we encountered in this project was that of efficiently clustering microbial genes based on co-abundance. This general approach has been proposed and implemented previously [13, 16], but existing implementations do not perform exhaustive searches for co-abundant genes because performing all pairwise comparisons of millions of genes in large microbiome datasets [17] is computationally intractable. To overcome this obstacle we took advantage of the Approximate Nearest Neighbor (ANN) heuristic, which is able to robustly identify candidate subsets of co-abundant genes without having to perform all pairwise comparisons [18, 19]. We implemented a Python package (“ann_linkage_clustering”) to perform exhaustive average linkage clustering using the cosine distance metric on any dataset containing gene abundance data across a set of samples. While this method is relatively computationally intensive, we were able to execute it in a reasonable amount of time using commodity “cloud” computational resources (e.g., 17 hours for a set of 5 million genes across 199 samples with a 256GB RAM node). While this clustering procedure is not expected to be deterministic, our experience has been that clusters are generally reproduced across replicates and we are actively studying the generalizability of gene clustering as a function of input data and clustering thresholds. In the ideal case this approach improves the precision of estimating gene-level abundance by combining data from multiple correlated observations, as well as reducing the number of hypotheses to test in an association study, while maintaining the interpretation advantages of distinct genetic elements (core genome, plasmid, virus, etc.).

We applied this novel approach to gene-level metagenomics to test for an association of the gut microbiome across two distinct human diseases: IBD and CRC. We selected these diseases because each has been studied by multiple groups who have collected stool samples and performed metagenomic WGS sequencing (Table S1) [2–8]. Because each of these previous studies used slightly different protocols for selecting patients, collecting samples, and performing sequencing, an integrated analysis of these datasets should serve to identify those signals in the microbiome which are most robust to the methodological and experimental confounders.

The CAGs identified in this project contained 2-23,856 genes, with the majority of genes found in CAGs ranging between 10 and 2,000 genes in size and containing the range of metabolic functions expected from complete and partial microbial genomes (Fig. S1). Visual inspection of the genes making up these CAGs also demonstrated the highly consistent patterns of abundance displayed by the genes which were ultimately grouped into these CAGs (Fig. 1). We also analyzed a published single-cell sequencing dataset from the stool microbiome [20] and found that genes from the same CAG were found in the same physical cell at 3-9X the rate expected by chance (Fig. S2). The size, functional content, and clear pattern of co-abundance displayed by the genes in this analysis suggest that the CAGs used for statistical analysis represent biological units that are meaningful reflections of the composition of the microbiome across multiple independent datasets.

**Figure 1.**
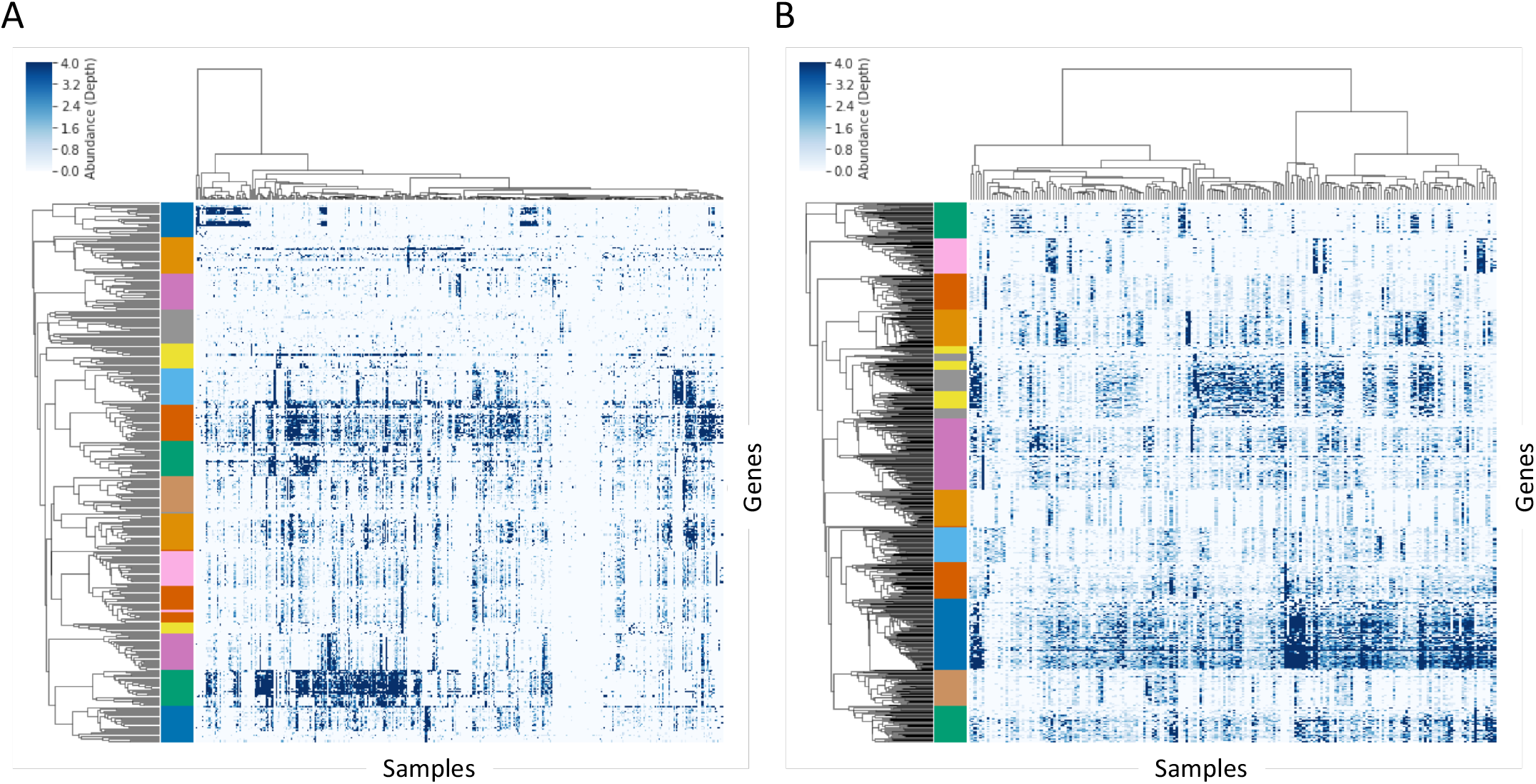
Patterns of gene-level co-abundance across all microbiome samples from a subset of CAGs. Each row represents a single microbial gene, each column represents a single biological sample, and pixel color reflects the gene’s relative abundance (sequencing depth) in the sample. A subset of CAGs and genes was randomly selected for display from the CRC datasets (A) and the IBD datasets (B). Unsupervised hierarchical clustering was used to group the rows and columns, and the left-hand color bar indicates the CAG assignment for each gene.

Our approach to the bioinformatic and statistical analysis was to select a single study for each disease as the “discovery” cohort, and to use that dataset to build a *de novo* catalog of microbial genes and identify CAGs. That gene catalog and CAG grouping generated from the discovery cohort was subsequently used to analyze the additional validation cohorts. Our statistical model was relatively straightforward and used random effects modeling to estimate the difference in the centered-log ratio of the relative abundance of each CAG in the samples from people with and without the disease state (accounting for multiple sampling of some individuals with random effects models). We chose to group together all participants with any form of the disease state, as the criteria for disease classification was not consistent across studies. In this discovery-validation approach, those CAGs which had a q-value of < 0.2 in the discovery cohort were subsequently tested in an additional “validation” cohort, and those CAGs which also had a q-value < 0.2 in that second step and the same direction of effect were considered to be associated with disease.

We found with this approach that the estimated coefficient of disease status in the set of CAGs associated with disease in the discovery cohort was significantly associated with the estimated coefficient in the validation cohort (Fig. 2A-B; CRC r=0.36 p<2E-16; IBD r=0.30 p<2E-16). Within the set of CAGs that were associated with disease in the discovery dataset, 44.0% and 97.2% were significantly associated in the validation dataset for CRC and IBD, respectively. When performing the same analysis with unclustered gene-level abundances (a single gene randomly selected from each CAG), we found a roughly 20-40% lower correlation between the estimated coefficient of disease status (Fig 2C-D) and a much lower validation rate of 9.8% and 76.0%, respectively. We believe that this evidence supports the proposed utility of CAGs for detecting reproducible biological associations of the microbiome with host disease. Furthermore, 24,502/36,871 CRC-associated CAGs had the same sign of the estimated coefficient in the validation cohort as in the discovery cohort (p < 1E-200, see Methods), and 28,629/31,895 IBD-associated CAGs had the same signed estimated coefficient (p < 1E-200). We further demonstrated the extent of this association by displaying the abundance of the most strongly associated CAGs across a total of 3 (IBD) or 4 (CRC) cohorts (Fig 2E-F), suggesting that this association is not limited to the cohorts selected for discovery and validation. Over and above the claim that the microbiome is associated with disease in both cohorts, we believe that these results indicate that a substantial number of elements of the microbiome that are associated with disease in a given discovery cohort will also be associated with disease in a corresponding validation cohort.

**Figure 2.**
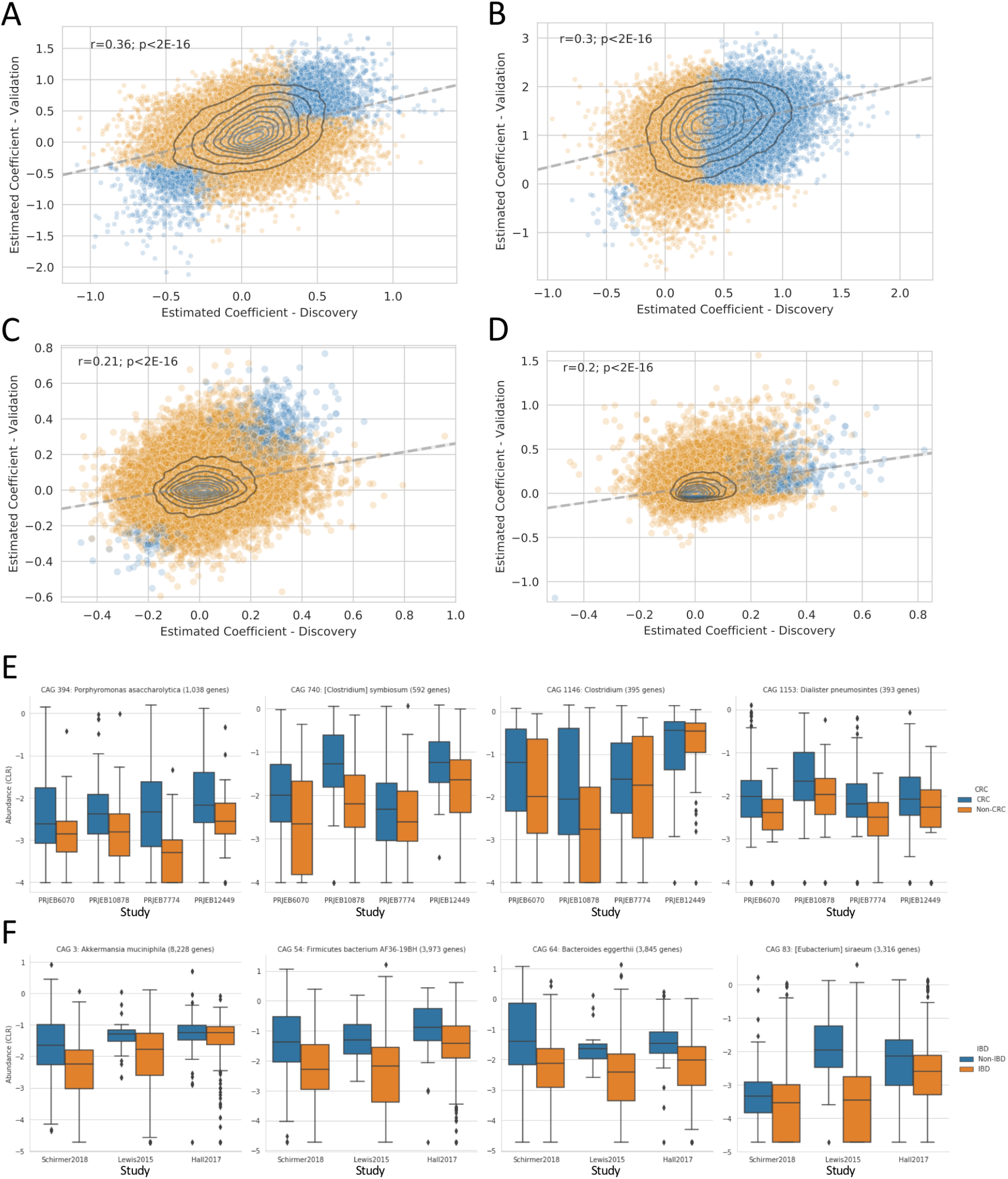
Reproducible association of CAG abundance with disease status for CRC (A, E) and IBD (B, F). The estimated coefficient plotted in A-D represents the log10 change in relative abundance associated with health (positive values) or disease (negative values), for each disease state. The estimated coefficient for the discovery dataset is on the horizontal axis, and the estimated coefficient for the validation dataset is on the vertical axis. The results from CAG-based analysis are shown in A-B, while the results calculated from unclustered gene-level abundances are shown in C-D. The abundance of four representative CAGs are shown in E-F across all available datasets, with colors indicating the health status associated with each sample.

The pattern of association for the IBD datasets was dominated by the 98.5% of CAGs which had a positive coefficient, indicating that they were more abundant in participants without IBD (Fig 2B). We therefore investigated the gene-level richness, finding a lower level of gene richness observed in IBD samples compared to healthy controls (Fig S3) [21], corroborating previous observations of lower alpha diversity in IBD [22, 23]. Without our use of the centered log-ratio to adjust for the compositional nature of these datasets the decreased abundance of a large fraction of the microbiome may have resulted in a spurious finding that the remainder had increased in abundance [24], but in fact we found that very few CAGs were consistently increased in abundance in IBD relative to the geometric mean of each sample. In addition to the decrease of overall gene richness, the lower number of CAGs found to be consistently enriched in IBD may also be due to an overall heterogeneity or ‘dispersion’ in the organisms which are positively associated with IBD across different people at a given point in time [14, 25]. However, there was a subset of CAGs which were consistently found to be more abundant in IBD, which may represent those bacteria which are able to thrive in the environment of the inflamed gut. Indeed, the taxonomic annotation of the genes in these CAGs is enriched for organisms which have been implicated in some previous studies of IBD and gut pathogens, including Enterobacteriaceae such as Escherichia/Shigella and Salmonella [3, 22, 23] which may exhibit some growth advantage in the context of either the increased oxygen content of the inflamed intestine or the antibiotics used in IBD treatment [9, 10]. Other organisms, such as *Ruminococcus gnavus*, were only enriched in IBD for a subset of genes (n=77), supporting the previous hypothesis of a strain-specific association with IBD [4]. There was also a set of KEGG annotations that were weakly but consistently enriched in this set of IBD-associated genes related to colonization and pathogenesis, such as fimbriae genes fimA (K07345) and fimD (K07347), iron transport (K02010), and putrescine transport (K02052; K11072; K11076).

The pattern of association for the CRC datasets was generally balanced between CAGs that were more abundant in healthy participants and those that were more abundant in disease (Fig 2A). Of the largest CAGs that were reproducibly associated with disease, those which were more abundant in healthy participants tended to be classified as Clostridia (via alignment to NCBI RefSeq), while those which were more abundant in participants with CRC were more taxonomically diverse (Fig 3A-B). Moreover, we found the functional annotations of the genes in those CAGs to be particularly interesting. There were four KEGG annotations that were significantly enriched in the set of CAGs found to be more abundant in CRC samples (Fisher’s exact test, Holm-Sidak alpha=0.01): 1) grdA (K10670) is involved in metabolism of glycine/sarcosine/betaine, and higher levels of glycine is a recognized hallmark of cancer cells [26, 27]; 2) oxyR (K04761) is a transcriptional regulator which regulates genes protecting from the biochemical damage induced by reactive oxygen species, of which markedly higher levels are associated with progressive tumors [28, 29]; 3) abgT (K12942) is a transporter responsible for uptake of p-aminobenzoyl-glutamate, and may also import other dipeptides [30]; and 4) afuA/fbpA (K02012) are transporters responsible for importing iron [31], which is likely to be more abundant in the gastrointestinal lumen of individuals with CRC due to bleeding. Three of these four annotated functions have clear links to the altered environment of the gut microbiome expected during CRC, and likely promote the growth of these organisms in that setting. It remains to be seen whether those organisms which are able to thrive in the CRC gut microbiome also contribute to progression of disease.

**Figure 3.**
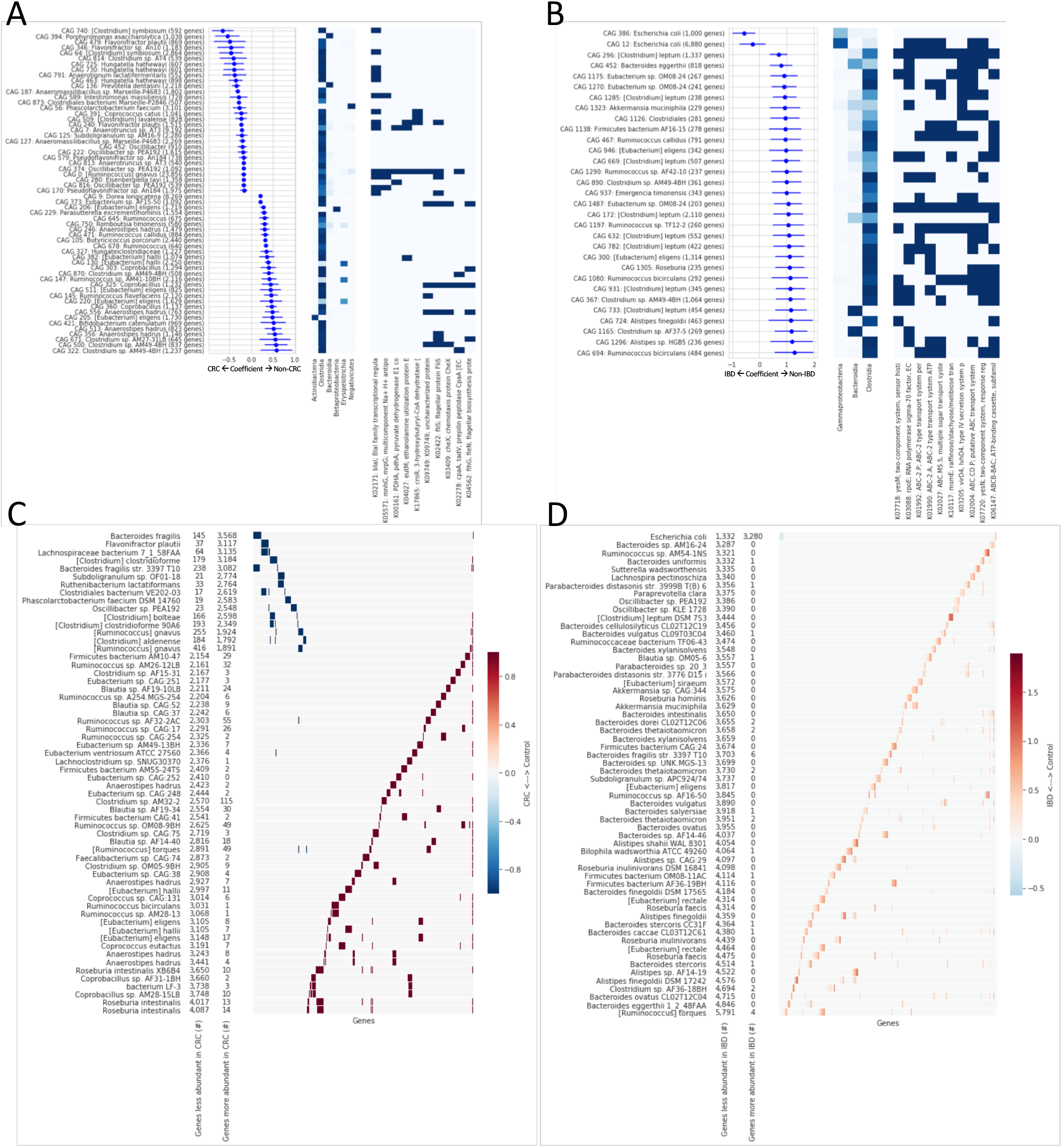
Association of individual microbes with CRC (A & C) and IBD (B & D). A & B show the estimated coefficient of abundance for individual CAGs with disease status (log10 mean and 90% confidence intervals, left panel), the taxonomic assignment (middle panel) and functional assignment (right panel) of genes within each of those CAGs. C & D show the number of genes from disease-associated CAGs that are found within bacterial genomes from NCBI RefSeq, showing both the total number of genes for each genome, as well as a heatmap showing which disease-associated genes are found in which genomes.

One advantage of a gene-based approach to metagenomic analysis is that any CAG of interest can be directly compared with the genomes of bacterial isolates in order to identify strains containing each gene. Of the set of genes that we identified as consistently associated with CRC and IBD, we found a number of strains containing large fractions of these genes (Fig 3C-D). We furthermore propose that this approach of aligning disease-associated genes to whole microbial genomes may be used to identify the members of any culture collection which are likely to have the largest effect in an experimental model of these human diseases.

## Conclusions

Having identified microbial protein-coding genes that are associated with CRC and IBD, we anticipate that other researchers may build on these findings in multiple ways. Researchers may compare this list of disease-associated genes to any genomes of interest in order to identify specific isolates and/or genes which may be perturbed in a controlled experimental setting to test the effect of microbes on host disease. Additionally, researchers may apply this general approach (quantification of CAGs from a *de novo* gene catalog) to their own metagenomic datasets in order to identify additional genes associated with any outcome of interest. While latter use-case may be implemented using the computational tools and associated Docker images described in the Methods, we are hoping to further support this methodological approach by developing reproducible analytical workflows that are more easily executed by the general microbiome research community.

By proposing an approach to the analysis of metagenomic data that produces consistent results across multiple heterogeneous datasets, we are addressing one of the most important challenges in metagenomics, namely, reproducibility. Our findings suggest that indeed co-abundant gene groups are a reproducible and biologically meaningful unit of analysis. In addition, microbial genes are a meaningful and useful unit of analysis because they can be linked to individual microbial genomes, taxonomic annotations, and predicted metabolic functionality. Using this approach, we identify a list of gene groups that are associated with human diseases in multiple cohorts, and we identify specific microbial isolates that contain these genes. The development of diagnostics or therapeutics based on this list of genes and genomes is left to future work.

## Methods

Datasets

### Gene-level metagenomic analysis pipeline

All microbiome WGS data was analyzed using a Docker-based workflow, with each individual step executed inside a Docker image. The workflow outlined below was executed independently for the set of samples from Schirmer, et al., as well as for the set of samples from Zeller, et al.

The sequence of analyses is as follows:

1. Each sample was individually downloaded from NCBI SRA with Entrez Direct

- <u>Docker image</u>: quay.io/fhcrc-microbiome/get_sra:v0.4
- <u>Code</u>: https://github.com/FredHutch/docker-sra
- <u>Wrapper script</u>: get_sra.py
- <u>Software version(s)</u>:

- sratoolkit.2.8.2-ubuntu64
- CMake3.11
- fastq-pair 4ae91b0d9074410753d376e5adfb2ddd090f7d85
2. Each sample was individually assembled with metaSPAdes

- <u>Docker image</u>: quay.io/fhcrc-microbiome/metaspades:v3.11.1--10
- <u>Code</u>: https://github.com/FredHutch/docker-metaspades
- <u>Wrapper script</u>: run_metaspades.py
- <u>Software version(s)</u>: SPAdes-3.11.1-Linux
3. Each sample’s metagenomic assembly was annotated using Prokka

- <u>Docker image</u>: quay.io/fhcrc-microbiome/metaspades:v3.11.1--8
- <u>Code</u>: https://github.com/FredHutch/docker-metaspades
- <u>Software version(s)</u>: Prokka v1.12; barrnap v0.9
- <u>Wrapper script</u>: run_prokka.py
4. The protein-coding sequences from all of the metagenomic assemblies for a given dataset were clustered at 90% amino acid identity using mmSeqs2 to create a set of non-redundant protein sequences

- <u>Docker image</u>: quay.io/fhcrc-microbiome/integrate-metagenomic-assemblies:v0.4
- <u>Code</u>: https://github.com/FredHutch/integrate-metagenomic-assemblies
- <u>Software version(s)</u>: biopython==1.70; MMseqs2 v2-23394
- <u>Wrapper script</u>: integrate_assemblies.py
5. Each sample was aligned against the non-redundant protein sequences using DIAMOND, with post-alignment filtering using FAMLI. The Docker image associated with this step includes both the DIAMOND aligner and the FAMLI filtering code.

- <u>Docker image</u>: quay.io/fhcrc-microbiome/famli:v1.1
- <u>Code</u>: https://github.com/FredHutch/famli
- <u>Software version(s)</u>: DIAMOND v0.9.10; famli==1.0
- <u>Wrapper script</u>: famli
- <u>Parameters</u>:

- min_qual = 30
- min_score = 20
- query_gencode = 11
6. The non-redundant protein sequences were functionally annotated via eggNOG-mapper

- <u>Docker image</u>: quay.io/fhcrc-microbiome/eggnog-mapper:v0.1
- <u>Code</u>: https://github.com/FredHutch/docker-eggnog-mapper
- <u>Software version(s)</u>: eggNOG-mapper = 1.0.3--py27_0
- <u>Wrapper script</u>: run_eggnog_mapper.py
7. The non-redundant protein sequences were analyzed via the taxonomic assignment functionality of DIAMOND (using NCBI’s RefSeq as the reference database)

- <u>Docker image</u>: quay.io/fhcrc-microbiome/famli:v1.3
- <u>Code</u>: https://github.com/FredHutch/famli
- <u>Software version(s)</u>: DIAMOND v0.9.22
- <u>Wrapper script</u>: diamond-tax.py
- <u>Parameters</u>: top_pct = 1
8. The non-redundant protein sequences were grouped into CAGs based on their abundance profile across the dataset.

- <u>Docker image</u>: quay.io/fhcrc-microbiome/find-cags:v0.11.1
- <u>Code</u>: https://github.com/FredHutch/find-cags
- <u>Software version(s)</u>: nmslib = 1.7.3.5
- <u>Wrapper script</u>: find-cags.py
- <u>Parameters</u>:

- min_samples = 10
- max_dist = 0.3
- normalization = sum
- Group the outputs of all previous steps into a single HDF file

- <u>Docker image</u>: quay.io/fhcrc-microbiome/experiment-collection:latest
- <u>Code</u>: https://github.com/FredHutch/minot-experiment-collection
- <u>Wrapper script</u>: make-experiment-collection.py

The validation datasets were analyzed by aligning the raw WGS reads against the non-redundant protein sequences generated from the relevant discovery dataset as described in Step 5 described above. The final HDF file creation step (9) includes the results of that quantification step for the validation datasets as well as the discovery datasets.

Given the difficulty of providing a workflow execution system that can be used effectively by a broad range of users, we have elected to provide all of the individual tools needed to run a complete analytical workflow, with public Docker images making up each individual step, instead of providing a complete workflow system that each user would need to customize for their own execution engine (Slurm, PBS, Kubernetes, AWS, GCP, Azure, etc.). This approach enables execution of the exact code that we used in this analysis in a platform-independent manner using the highest standard of reproducibility (Docker containers).

Our implementation of the analytical workflow described above relied upon the Amazon Web Service and its Batch API, which allows users to submit individual jobs for analysis using utilities from the boto3 library in Python. While this implementation does not represent a complete workflow management system, the code used for this execution is available at https://github.com/FredHutch/aws-batch-helpers/ in the batch_helpers/batch_task_manager.py module.

### Grouping genes by co-abundance

We did not find any public tools for grouping genes by co-abundance that were appropriate to the scale of our datasets. To implement our own approach for finding CAGs, we utilized the Non-Metric Space Library (‘nmslib’, https://pypi.org/project/nmslib/) which implements the Approximate Nearest Neighbor (ANN) algorithm [18, 19] and obviates the need for calculating the all-by-all distance matrix typically used by clustering algorithms. The abundance matrix used for clustering was created by calculating the depth of sequencing for each individual gene within each sample and normalizing for total sequencing depth. The distance metric used to quantify the dissimilarity of individual genes was the cosine distance. Gene clusters were identified iteratively by average linkage clustering and a fixed cophenetic distance threshold. The ANN algorithm was used to identify subsets of genes which were likely to be highly co-abundant, and which could be clustered independently of the whole. The code executed for this analysis, as well as a Docker image containing all required dependencies, can be found in the summary of the complete analysis workflow (Step 8).

### Correlating CAGs with health status

#### CAG discovery

For every CAG in the validation dataset, we tested the null hypothesis that the mean difference in CLR abundance between patients with and without disease was zero using the general linear model framework. Datasets with repeated measurements on subjects were modelled using a linear mixed effects model with subject as a random effect. We employed the centered-log ratio to address the compositionality and range constraint of the gene relative abundances, and it is consistent with the choice to group genes based on cosine distance. Using the `qvalue` R package (v2.8.0), we calculate the q-values for each CAG. Our set of “discovered CAGs” for validation is the set of CAGs with calculated q-value of 0.2 or less. These are the CAGs that would be considered statistically significant while controlling the FDR at 20%.

#### CAG validation

*For only the discovered CAGs*, we tested the null hypothesis that the mean difference in CLR abundance between patients with and without disease was zero in the validation datasets. Our “validated CAGs” are the CAGs in this set with calculated q-values of 0.2 or less, and that have an estimated difference in abundance between disease status groups of the same sign as the estimated difference in the discovery dataset.

#### The probability of validating discovered CAGs

To calculate the probability of validating C_2_ or more out of C_1_ CAGs under the global null hypothesis of no association between disease status and any CAG’s abundance, we bounded the p-value for validating discovered CAGs in the following way. Let X be the number of CAGs with q-values less than 0.2 for the validation data and with an estimated difference in CLR abundance across disease groups of the same sign in the validation and discovery datasets, and Y be the number of CAGs with an estimated difference in CLR abundance across disease groups of the same sign in the validation and discovery datasets. Since under the null the test statistics are approximately Normal(0,1)-distributed,

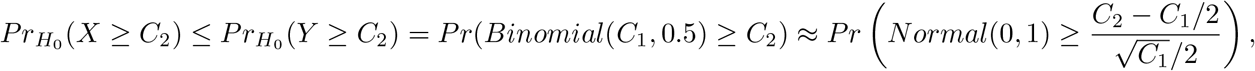

giving us a conservative p-value for the global null of no association.

### Aligning protein-coding genes against RefSeq genomes

The alignment of individual protein-coding genes against the RefSeq collection of genomes in NCBI was executed using the Docker image hosted at quay.io/fhcrc-microbiome/docker-diamond:v0.9.23—0 and built using the Dockerfile hosted at https://github.com/FredHutch/docker-diamond, running DIAMOND v0.9.23. The complete list of Prokaryotic RefSeq genomes was downloaded from https://www.ncbi.nlm.nih.gov/genome/browse#!/prokaryotes/ and the query proteins were aligned via DIAMOND against the annotated protein-coding sequences from each genome individually. We implemented this analysis on the Amazon Web Service using the Batch API for execution and resource management.

### Quantification of co-abundant genes in uncultured single cells

Datasets from published single-cell sequencing microbiome experiments [20] were downloaded and split by 10X barcode (each corresponding to a single cell). The WGS data for each single cell was aligned against each reference gene catalog (for the CRC and IBD datasets) and filtered with FAMLI as described in workflow step 5, above. The result of this analysis was a count of the number of genes that were found in the same cell as another gene that is also part of the same CAG. As a comparison, we calculated the number of such genes that would be found with a randomly permuted set of CAG assignments.

## Declarations

### Ethics approval and consent to participate

Not applicable

### Consent for publication

Not applicable

### Availability of data and material

The data produced in this analysis is available on the Synapse platform at https://www.synapse.org/#!Synapse:syn15623121 (doi:10.7303/syn15623121). The Synapse project will be made fully public upon acceptance for publication. The repository includes documentation describing the organization and formatting of relevant data files and includes all of the outputs from the bioinformatic pipeline used for gene-level metagenomic analysis, as well as the Jupyter notebooks used to analyze those datasets and produce the figures and tables presented here.

### Authors’ contributions

SM developed the novel CAG identification method and performed all primary data analysis; AW developed the statistical analysis and discovery-validation framework; both authors contributed to figure generation and writing the manuscript.

## Acknowledgements

The authors are grateful for support from the Fred Hutch Microbiome Research Initiative (SM), start-up funds from the Department of Biostatistics at the University of Washington (AW), and valuable intellectual contribution and support from Dr. Jonathan L. Golob.

## Supplementary Figures

**Supplementary Figure S1.**
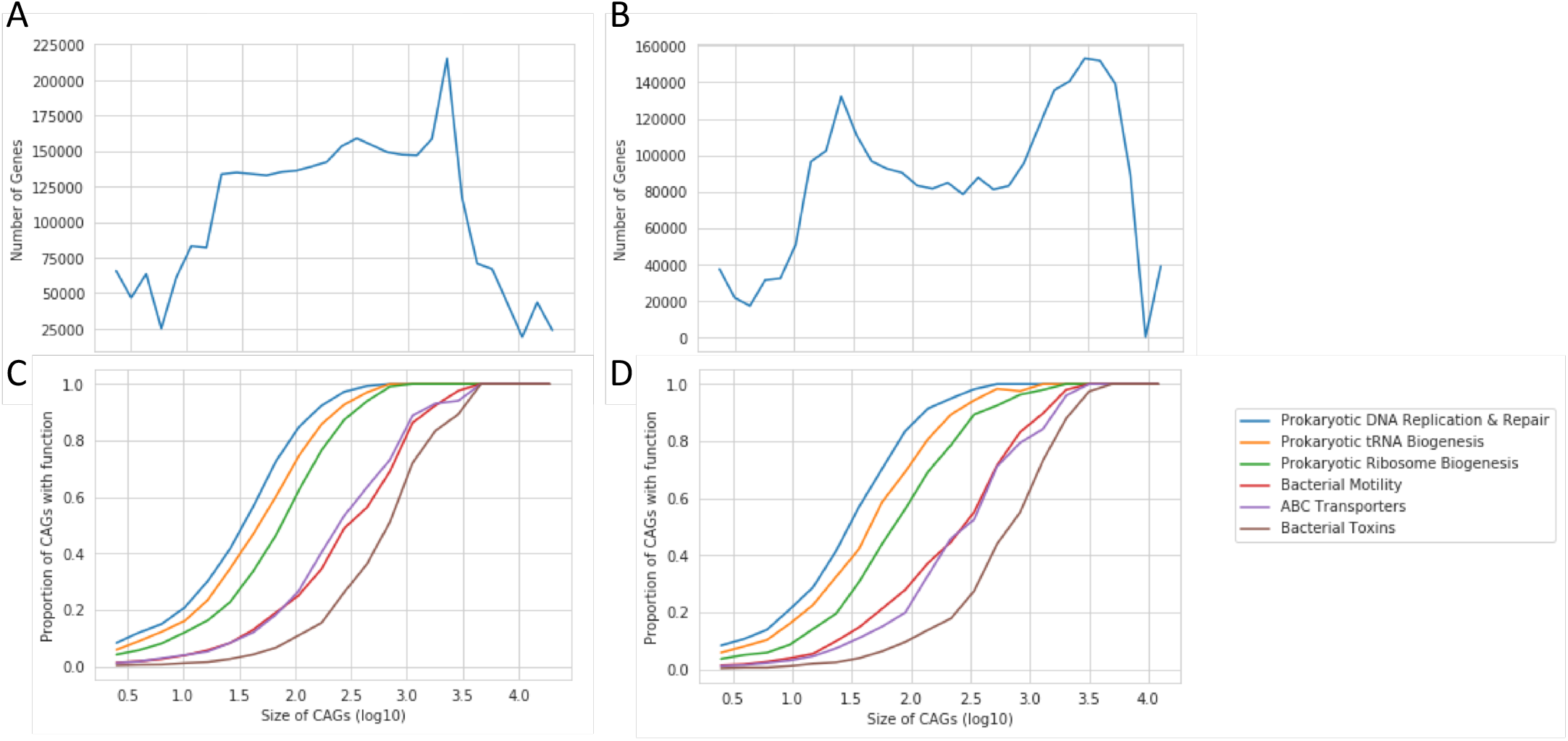
The distribution of CAG size (genes per CAG; A & B) and the functional annotation of genes in CAGs is shown by CAG size (C & D). Each gene can be annotated with a range of biological functions, and the proportion of CAGs of a given size containing at least one functional annotation is shown (C & D). The CAGs generated from the CRC datasets are shown in A & C, while the CAGs generated from the IBD datasets are shown in B & D. The horizontal axis is shared between panels A & C, as well as B & D.

**Supplementary Figure S2.**
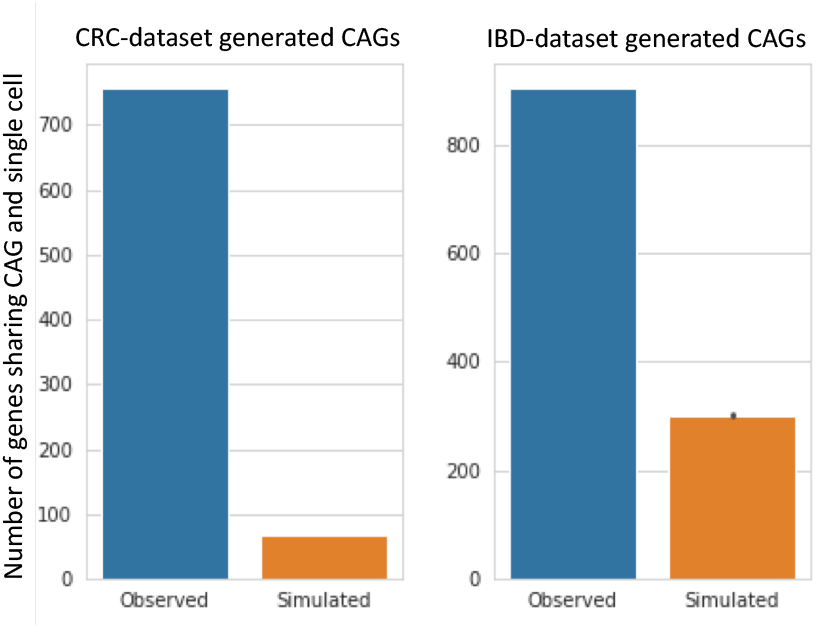
Single-cell microbiome datasets were analyzed using the gene catalogs and CAG groupings from the CRC and IBD datasets. Co-occurrence was measured as the number of genes that were found in the same cell with another gene from the same CAG. Simulations were performed by random permutation, with 1,000 replicates. Orange bars show mean and standard deviation.

**Supplementary Figure S3.**
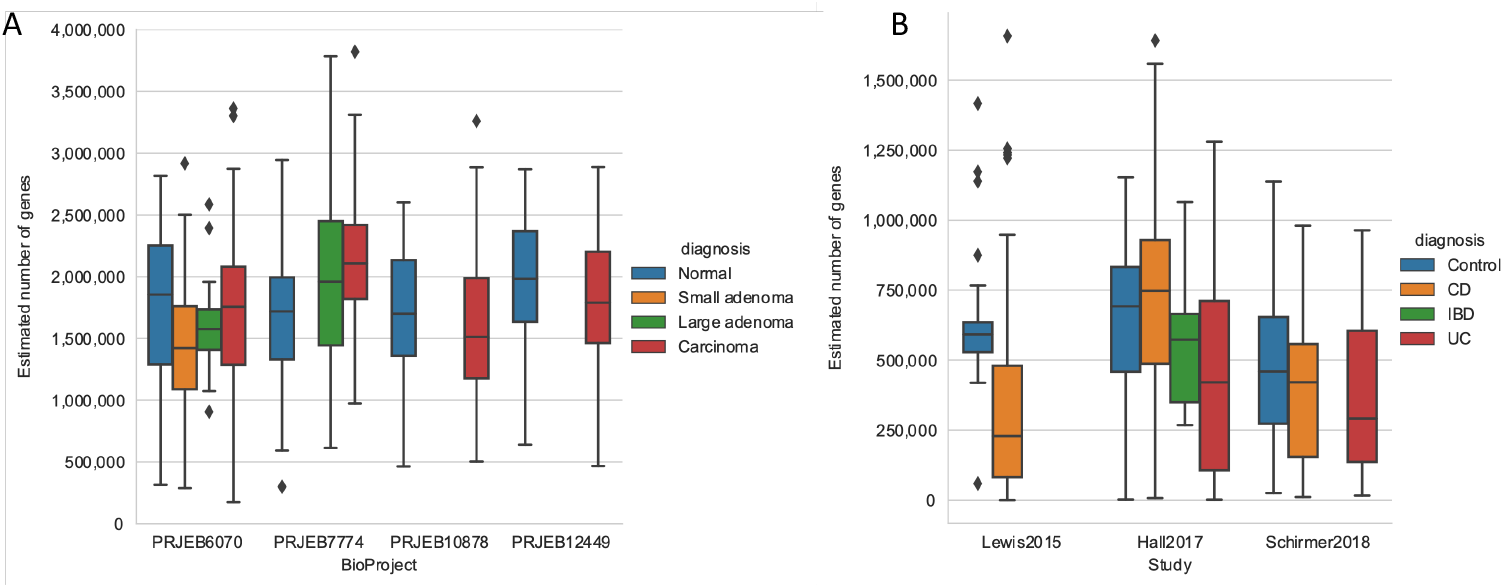
Alpha diversity by diagnosis across cohorts. The number of total genes in each sample was estimated with breakaway for both the CRC (A) and IBD (B) cohorts.

## Supplementary Tables

**Table S1.** Published datasets analyzed in this study.

| Group | Used For | Name | NCBI BioProject |
| --- | --- | --- | --- |
| IBD | Discovery | Schirmer, et al. [2] | PRJNA389280 |
| IBD | Validation | Lewis, et al. [3] | SRP057027 |
| IBD | Validation | Hall, et al. [4] | PRJNA385949 |
| CRC | Discovery | Zeller, et al. [5] | PRJEB6070 |
| CRC | Validation | Feng, et al. [8] | PRJEB7774 |
| CRC | Validation | Yu, et al. [7] | PRJEB10878 |
| CRC | Validation | Vogtmann, et al. [6] | PRJEB12449 |

**Supplementary Table S2.** Description of genes associated with CRC, including the CAG grouping, correlation coefficient, taxonomic annotation, and functional annotation. Public repository URL: https://www.synapse.org/#!Synapse:syn17104367

**Supplementary Table S3.** Description of genes associated with IBD, including the CAG grouping, correlation coefficient, taxonomic annotation, and functional annotation. Public repository URL: https://www.synapse.org/#!Synapse:syn17104250

